# Coronavirus RNA synthesis takes place within membrane-bound sites

**DOI:** 10.1101/2021.11.04.467246

**Authors:** Nicole Doyle, Jennifer Simpson, Philippa C Hawes, Helena J Maier

## Abstract

Infectious bronchitis virus (IBV), a gammacoronavirus, is an economically important virus to the poultry industry as well as a significant welfare issue for chickens. As for all positive strand RNA viruses, IBV infection causes rearrangements of the host cell intracellular membranes to form replication organelles. Replication organelle formation is a highly conserved and vital step in the viral life cycle. Here, we investigate the localization of viral RNA synthesis and the link with replication organelles in host cells. We have shown that sites of viral RNA synthesis and virus-related dsRNA are associated with one another and, significantly, that they are located within a membrane-bound compartment within the cell. We have also shown that some viral RNA produced early in infection remains within these membranes throughout infection. Importantly, we demonstrate conservation across all four coronavirus genera, including SARS-CoV-2. Under-standing more about the replication of these viruses is imperative in order to effectively find ways to control them.

## Introduction

Coronaviruses (CoVs) are an important family of positive strand RNA (+RNA) viruses with a wide host range. In humans, some strains of these viruses such as the human coronavirus (HCoV) 229E can cause the common cold, however there are now three CoVs that are more pathogenic and cause higher fatality rates. Until 2019, severe acute respiratory syndrome coronavirus (SARS-CoV) and Middle East respiratory syndrome coronavirus (MERS-CoV), which were initially isolated in China and Saudi Arabia, respectively [1, 2] were the most well-known zoonotic viruses in the coronavirus family. However, the emergence of SARS-CoV-2, causing global mortalities (over 4 million people at the time of writing [3]) and shut-down of normal life has brought a focus on the danger of zoonotic viruses, and in particular this virus family [4]. Apart from human viruses, coronaviruses also cause disease in a range of animal species. Several of these viruses are of economic importance as they cause infections and loss of income in the global agriculture industry. These include viruses such as porcine epidemic diarrhea virus (PEDV), porcine deltacoronavirus (PDCoV), bovine coronavirus (BCoV) and avian infectious bronchitis virus (IBV). As has recently been highlighted, studying these viruses is of vital importance. Much work on the CoV family over recent years has been focused on understanding how they interact with the host cell, including a key part of their life cycle, the induction of membrane rearrangements or replication organelles (ROs).

As obligate intracellular parasites, viruses rely on their host cells to provide not only a site for replication, but much of the cellular machinery required to produce new virus particles. All +RNA viruses induce the rearrangement of intracellular membranes to form ROs [5–7]. These viral ROs have been shown to be the site of viral RNA synthesis [8, 9], however it is probable that they also offer other benefits to the virus. The ROs are likely to provide a way to shield replicative intermediates which would otherwise be recognized by cellular defenses and spark an innate immune response [10, 11]. It has also recently been shown that ROs could be a site of local ATP production for the energy-intensive process of RNA synthesis which takes place there [12]. The structures formed vary between virus families but do share similarities in structures such as convoluted membranes, double membrane vesicles (DMVs) and spherules [13, 14]. Several viruses, such as toroviruses [15], hepatitis C virus (HCV; [16]) and picornaviruses such as foot and mouth disease virus (FMDV; [17]) induce the formation of tubules or paired membranes as well as single-membrane vesicles, DMVs or multi-lamellar vesicles. In enterovirus infections single-membrane tubules transform into DMVs and multilamellar vesicles over the course of infection [18, 19], using endoplasmic reticulum (ER) and Golgi membranes to initiate the formation of these structures [20]. A different structure induced by many other +RNA viruses are spherules or invaginated vesicles. These structures are smaller than DMVs and are pinched out from various intracellular membranes but because they remain bound to that membrane, they possess a channel which connects their interior to the cytoplasm. Viruses that induce spherule formation include flaviviruses [21–23], nodaviruses [24], bromoviruses [25] and alphaviruses such as Semliki Forest virus (SFV). In the case of SFV ROs, the spherules and the cytopathic vacuoles which contain them have been shown to be the site of viral RNA synthesis [26–28]. For flock house nodavirus (FHV), spherules have been found to form from the mitochondrial membrane [29] and these structures have also been shown to be the site of viral RNA synthesis, containing bundles of dsRNA [30, 31].

The possibility of antiviral therapies targeting the ROs has meant an increased attraction for understanding this part of the viral life cycle in recent years. Until very recently, the structures of ROs formed by the CoV family were split into the alpha- and beta-CoVs and the gamma- and delta-CoVs. The alpha- and beta-CoVs were known to induce convoluted membranes and DMVs [9, 32–35], and while DMVs are also produced by gamma- and delta-CoVs, these were seen alongside areas of tightly paired ER membranes called zippered ER (zER) and small double membrane spherules which, in IBV infected cells, have been shown to remain tethered via an open neck to the zER [36, 37]. However, it has recently been shown that alpha- and beta-CoVs also induce spherules, although with some morphological differences. They are formed to a lesser extent, and while some spherules appear to be connected to the convoluted membrane, many more were seen as sealed structures. While some sealed spherule structures were seen in IBV infection in the later study, this was to a lesser extent [8]. The significance of these differences between structures is not yet clear and is the source of further investigation.

As ROs have long been purported to provide a site for viral RNA synthesis, a key point for investigation has been in understanding exactly where the sites of viral replication are within the cell, and the involvement of ROs in the process. For SARS-CoV, it has been shown using biochemical methods *in vitro* that sites of viral RNA synthesis are protected by membranes, possibly within DMVs [38], and dsRNA signal was shown to be associated with DMVs and CM using immunostaining [39]. SARS-CoV-2 DMVs have also recently been shown to contain RNA filaments consistent with the size of dsRNA [40]. It is assumed that DMVs in mouse coronavirus (MHV) play a role in viral RNA synthesis based on the fact that mutations in one of the non-structural proteins, nsp4, affects both DMV formation and viral RNA synthesis [41, 42]. DMVs were known to be necessary for viral replication [43] and while DMVs have long been associated with sites of viral RNA synthesis [9], this theory was always hampered by the fact that DMVs were shown to be closed compartments with no way for newly synthesized viral RNA to escape for packaging and egress. However, DMVs have now been shown to be a site of viral RNA synthesis [8] and further recent findings demonstrated a small number of transient pores in the membranes of MHV DMVs, allowing egress of newly synthesized viral RNA [44]. The role of spherules remains to be elucidated.

The gammacoronavirus, IBV is a virus of economic importance to the global poultry industry, causing a highly contagious respiratory disease of chickens and other poultry. Infection results in reduced quantity of eggs and reduced quality of eggs and meat as well as impacting on animal welfare. Recent work has elucidated the role of ROs in viral RNA synthesis and suggested a pathway for nascent RNA to leave the DMV via a molecular pore [44]. However, conclusive proof of viral RNA synthesis occurring within the DMV was still lacking and the possibility remained that synthesis was taking place on the outer surface of DMVs. Here, we show that IBV viral RNA synthesis takes place within membrane-bound compartments, and we demonstrate that this is conserved across all four genera of the CoV family.

## Materials and Methods

### Cells and viruses

Avian DF1 cells (LGC Standards Ltd.) were maintained in DMEM (Sigma Aldrich, Gillingham, UK) supplemented with 10% FBS (Sigma Aldrich) at 37 °C 5% CO_2_. IBV (strain BeauR [45]) infections were carried out in 1xBES medium (MEM, 0.3% tryptose phosphate broth, 0.2% bovine serum albumin, 20 mM N,N-Bis(2-hydroxyethyl)-2-aminoethanesulfonic acid (BES), 0.21% sodium bicarbonate, 2 mM L-glutamine, 250 U/mL nystatin, 100 U/mL penicillin, and 100 U/mL streptomycin). Huh7 cells (ATCC) were maintained in DMEM supplemented with 10% FBS. Human coronavirus 229E (HCoV 299E (UK Health Security Agency) infections were carried out in DMEM + 2% FBS. VeroE6 cells (LGC Standards Ltd.) were maintained in DMEM supplemented with 10% FBS. SARS-CoV-2 (strain hCov-19/England/02/2020, kindly provided by Prof Miles Carroll, UK Health Security Agency) infections were carried out in DMEM +2% FBS. Porcine LLC-PK1 cells (ATCC CL-101 [46]) were maintained in DMEM supplemented with 10% FBS. PDCoV (strain OH-FD22, kindly provided by Prof Linda Saif, Ohio State University, [47, 48]), infection was carried out in EMEM + 1% HEPES, 1% non-essential amino acids, 1% antibiotic/antimycotic and 0.25 μg/mL trypsin. In all cases, cells were inoculated with virus for 1 h, after which time the inoculum was replaced with infection media specific to each virus. Cells were then incubated until specified timepoints.

### Labeling of nascent viral RNA with bromouridine

Cells seeded onto glass coverslips were infected as in 2.1. Cells were then treated with 2 mM (or 4 mM for HCoV 229E) bromouridine (BrU; Sigma Aldrich) and 15 μM actinomycin D (ActD; Sigma Aldrich) at 30 min prior to each timepoint. Cells were washed in PBS, fixed in RNase-free paraformaldehyde (pfm) at each timepoint, then labeled as in 2.3.

For pulse chase experiments, cells were pulsed with BrU and ActD for 1 h at concentrations previously used. After this time, control cells were fixed in RNase-free pfm, while the rest were chased with 50 μM uridine (Sigma Aldrich) to out-compete the labeled BrU. Cells were incubated in uridine and ActD until 24 hpi then fixed as above and labeled as in 2.3.

### Immunofluorescence labeling

Cells seeded onto glass coverslips were mock-infected or infected as laid out in 2.1. and 2.2. At each timepoint, cells were washed in PBS then fixed for 15 min in 4%pfm in PBS at room temperature. Cells were permeabilized in 0.1% Triton X-100 in PBS for 15 min, or 0.25% digitonin (BioVision Inc) in PBS for 10 min then incubated in blocking buffer (0.1% fish gelatin [Sigma Aldrich] in PBS) for 1 h. Primary antibodies specific for proteins of interest (Table 1) were diluted in blocking buffer and incubated on cells for 1 h. After washing, Alexa Fluor secondary antibody (Invitrogen) in blocking buffer was incubated on cells for 1 h, followed by washing, labeling of nuclei using 4’,6-diamidino-2-phenylindole (DAPI; Sigma Aldrich) or ToPro3 (ThermoFisher) and mounting onto glass slides with Vectashield (Vector Labs, Peterborough).

**Table 1.**
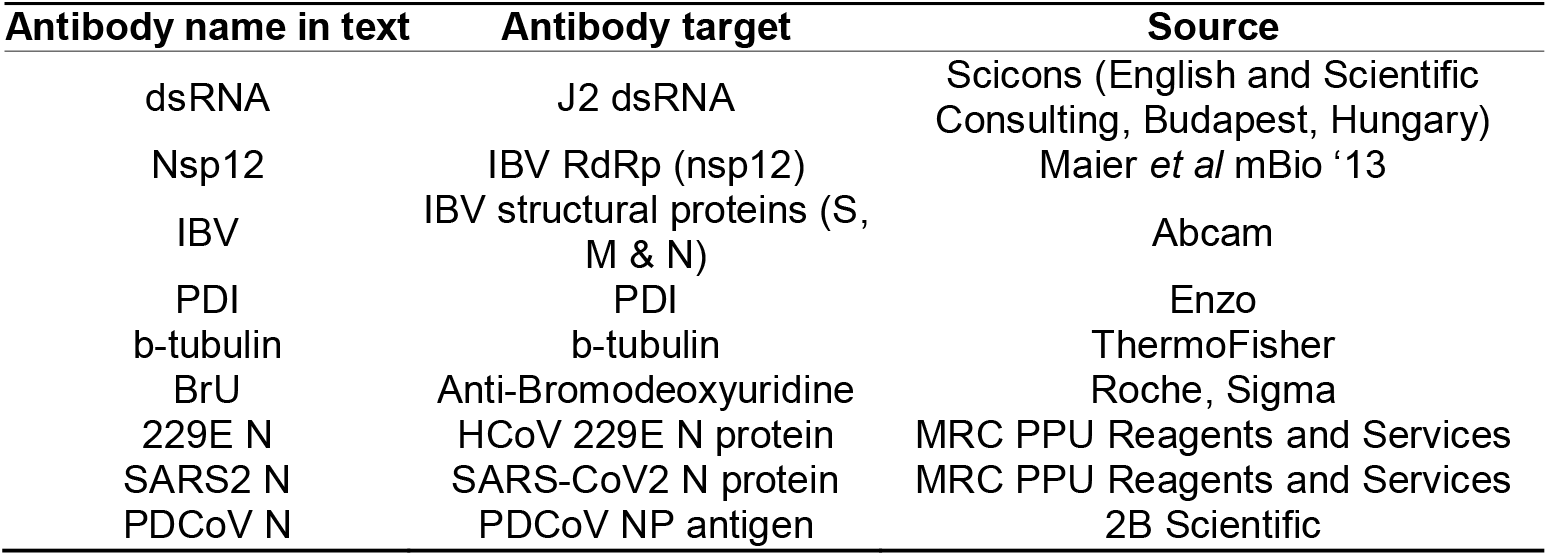
Table showing antibodies used in this paper

| Antibody name in text | Antibody target | Source |
| --- | --- | --- |
| dsRNA | J2 dsRNA | Scicons (English and Scientific Consulting, Budapest, Hungary) |
| Nsp12 | IBV RdRp (nsp12) | Maier *et al* mBio '13 |
| IBV | IBV structural proteins (S, M & N) | Abcam |
| PDI | PDI | Enzo |
| b-tubulin | b-tubulin | ThermoFisher |
| BrU | Anti-Bromodeoxyuridine | Roche, Sigma |
| 229E N | HCoV 229E N protein | MRC PPU Reagents and Services |
| SARS2 N | SARS-CoV2 N protein | MRC PPU Reagents and Services |
| PDCoV N | PDCoV NP antigen | 2B Scientific |

For labeling of BrU samples, in order to prevent loss of the BrU signal, immunofluorescence (IF) labeling was promptly carried out in an RNase-free environment in the presence of RNasin at 0.133 U/ml (Promega, Southampton; [49]).

Cells were visualized using a Leica CLSM SP5, SP8 or Stellaris 5 microscope (Leica Microsystems, Milton Keynes, UK). Super-resolution (stimulated emission depletion (STED)) microscopy was performed using a Leica TCS SP8 STED 3X microscope with inverted stand. STED Images were deconvolved using Huygens Professional software 18.10 (Scientific Volume Imaging, Netherlands). Figures were assembled using Adobe Photoshop.

## Results

### DsRNA is found in membrane-bound compartments

We have shown previously over the course of the IBV life cycle that nsp12, the viral RNA-dependent RNA polymerase (RdRp), does not colocalize with dsRNA [37]. The role of dsRNA is still a source of interest, but it is most likely to be a replicative intermediate, formed during the replication of the viral genome. Interestingly, dsRNA has recently been shown to be largely negative-sense viral RNA [50]. It has been shown in SARS-CoV that dsRNA is found within DMVs, which are quite likely to store it, hidden from detection by intracellular pattern recognition receptors [39]. The formation of dsRNA is a well conserved step across +RNA virus families so we sought to understand whether dsRNA is also protected within membrane-bound compartments in IBV infection using different permeabilization agents. Digitonin is a weak permeabilizing agent, which can be used at low concentrations to selectively permeabilize the plasma membrane but not intracellular membranes, while Triton X-100 (TX100) permeabilizes all cellular membranes. Mock-infected DF1 cells were permeabilized with TX100 (Fig 1, top) or digitonin (Fig 1, bottom), then labeled with antibodies specific for PDI and tubulin. Tubulin proteins are found in the cytoplasm, so tubulin labeling was visible upon either TX100 or digitonin permeabilization. In contrast, labeling of PDI, the intra-lumenal ER protein is lost with digitonin treatment as the antibody is prevented from accessing its target. DF1 cells infected with IBV and fixed at timepoints during the infection cycle as indicated (Fig 1) were similarly processed. On the top row, cells permeabilized with TX100 clearly show both nsp12 (green) and dsRNA (red) labeling, with the dsRNA labeling increasing markedly over the course of infection. However, when cells were permeabilized with digitonin (Fig 1, bottom row), dsRNA was no longer visible in the cells, indicating that it is held within an intracellular membrane and therefore inaccessible to the antibody. In comparison, nsp12 staining was unaffected when cells were permeabilized with digitonin (Fig 1), indicating that it is not contained within a membrane but is possibly free in the cytoplasm or on the cytoplasmic face of a membrane.

**Figure 1.**
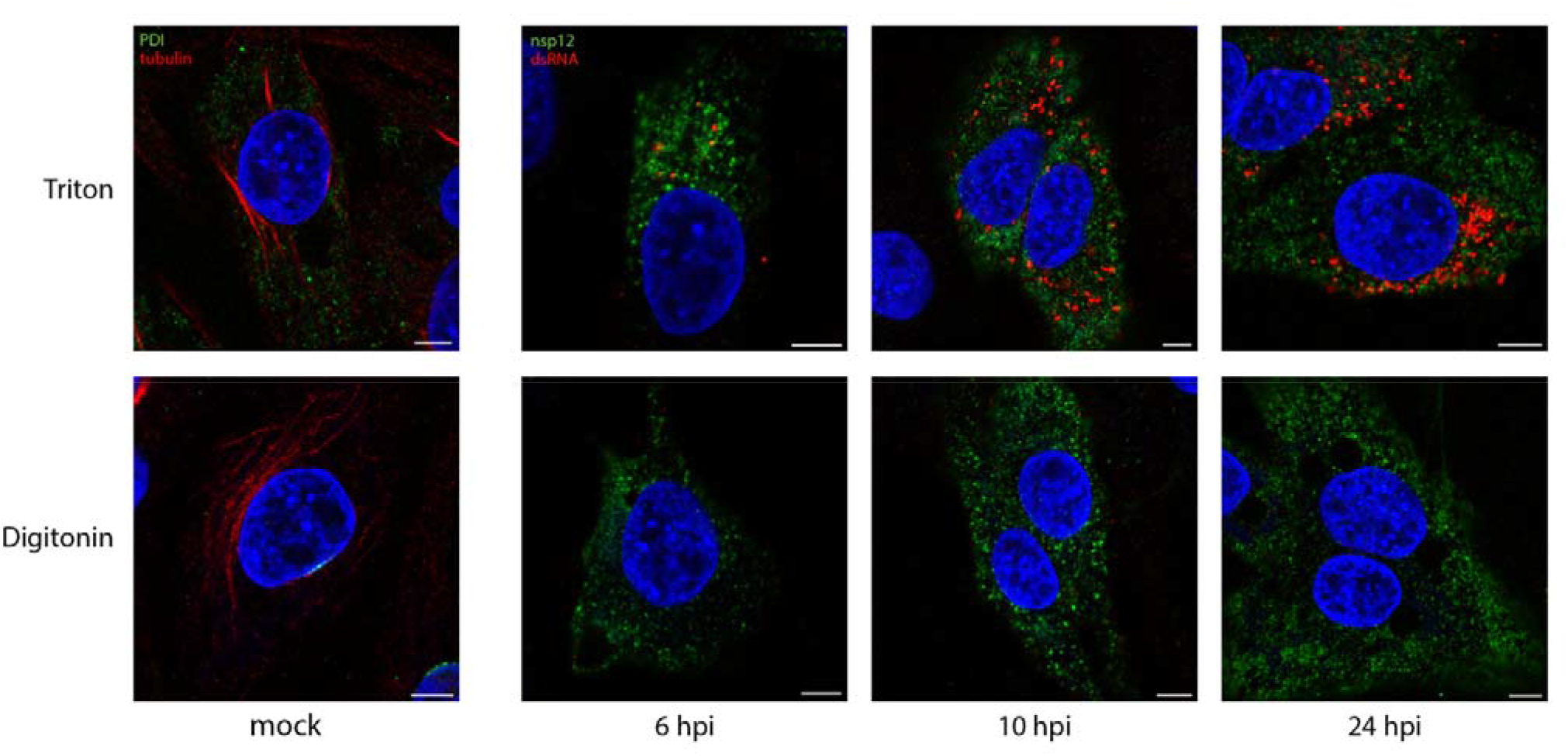
DsRNA is contained within a membrane-bound compartment whilst nsp12 is exposed to the cytoplasm. DF1 cells were infected with IBV and fixed at the indicated times post infection. Cells were permeabilized with either Triton X-100 (all membranes permeabilized; top row), or digitonin (plasma membrane permeabilized only; bottom row). Cells were labeled with dsRNA (red) and nsp12 (green), or for the mock control (first column), tubulin (red) and PDI (green). Nuclei are labeled blue with DAPI (blue). Scale bars represent 5 μm.

### DsRNA is closely associated with sites of viral RNA synthesis

Knowing that dsRNA was likely within the DMVs, we sought to understand how it associated with sites of viral RNA synthesis. To do this, the uridine analog bromouridine (BrU) was incorporated into the nascent viral RNA. DF1 cells were infected with IBV then incubated in media containing BrU for 30 min prior to fixation to provide a snapshot of RNA synthesis at that timepoint. The cellular transcription inhibitor, Actinomycin D (ActD) was used to selectively inhibit cellular transcription to allow visualization of sites of active viral RNA synthesis (Fig S1). Starting at 4 hpi, sites of viral RNA synthesis (Fig 2, green) were detected as small foci localized in the perinuclear region. As infection progressed, these sites of viral RNA synthesis increased in size and distribution around the cell, although more often than not retaining their perinuclear distribution. A similar pattern was observed for both nsp12 and dsRNA labeling (red), however while nsp12 labeling was generally diffuse over the cytoplasm, dsRNA labeling remained punctate and tended to accumulate in perinuclear regions, especially earlier in infection (Fig 2b). Over the course of infection with IBV, nsp12 did not colocalize with BrU (Fig 2a). This was surprising as we would expect the viral RdRp to be at sites of viral RNA synthesis, however no colocalization could be found even when analyzed in 3D (Fig S2 and Video S3) or using super resolution microscopy (Fig 2c). In contrast, dsRNA, appeared to exhibit a low level of colocalization or overlap with BrU signal at earlier timepoints, particularly in larger foci and as infection progressed, the overlap appeared to increase (Fig 2b). Using super-resolution microscopy, we confirmed that rather than tightly colocalizing, dsRNA and BrU signal were instead in close association with each other (Fig 2c). These results show that dsRNA, while not appearing to be completely colocalized with BrU-labeled nascent RNA, is closely associated with these sites of viral RNA synthesis.

**Figure 2.**
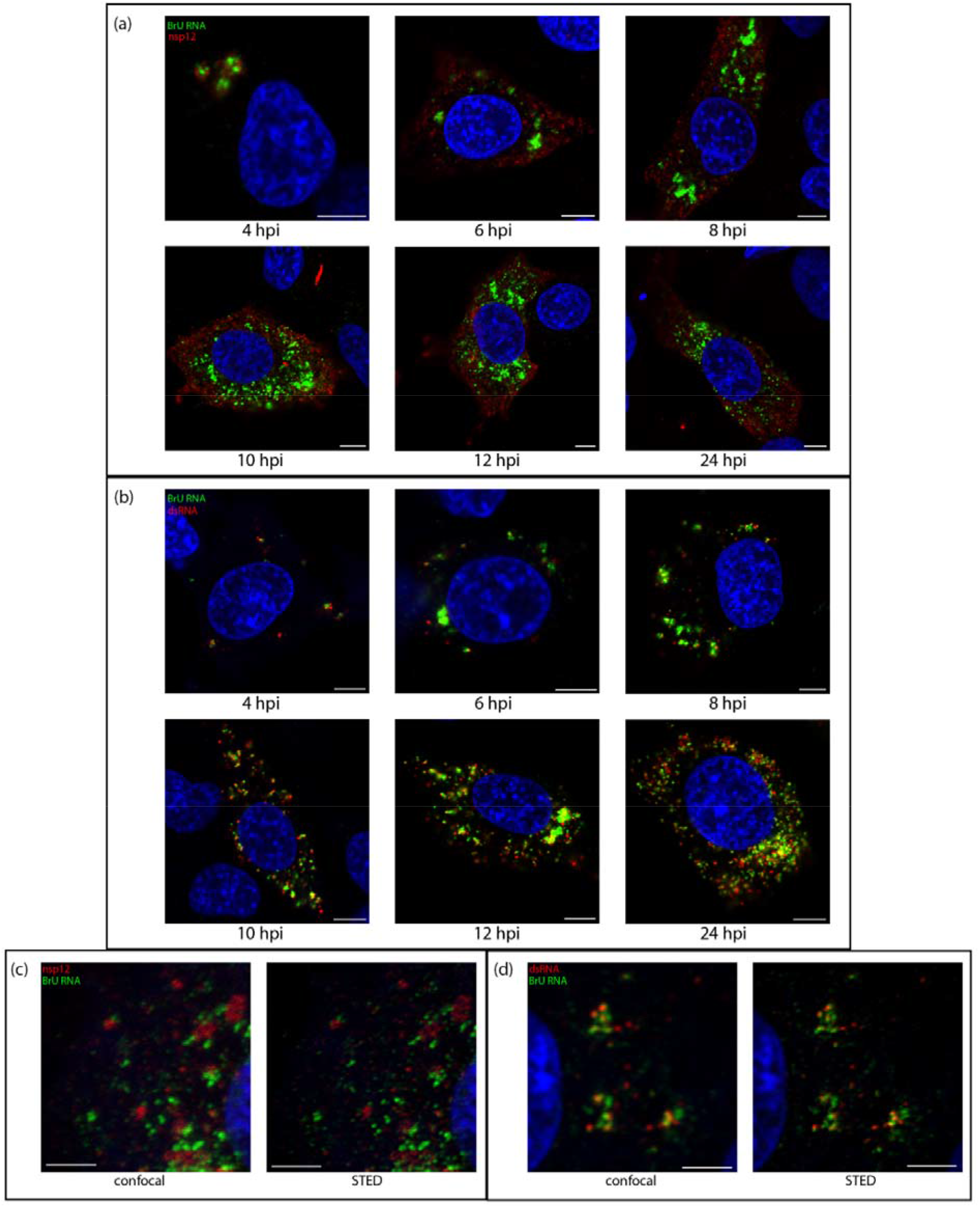
Sites of viral RNA synthesis are associated with dsRNA but do not colocalize with nsp12. DF1 cells were infected with IBV and 30 mins prior to fixation treated with BrU and ActD. Cells were fixed at the indicated time points post infection and labeled with antibodies against BrU (green) and a) nsp12 (red) or b) dsRNA (red). Nuclei are labeled with DAPI (blue). Scale bars represent 5 μm. (c & d) Cells were treated as in (a & b), fixed at 10 hpi and labeled for BrU (green) and c) nsp12 or d) dsRNA (red) and ToPro3 (blue) for the nuclei. A confocal image was captured followed by a super-resolution images which was captured using a STED microscope, then deconvolved. Scale bars represent 3 μm.

### Sites of viral RNA synthesis are membrane-protected

Snijder *et al* [8] showed that sites of viral RNA synthesis are associated strongly with ROs, in particular they showed they were more strongly associated with DMVs. While they subsequently showed that DMVs have a pore which would allow for the egress of newly synthesized viral RNA, the question of pinpointing nascent viral RNA within these structures remained [44]. Here it has been demonstrated that sites of IBV viral RNA synthesis and dsRNA are closely associated rather than precisely colocalized. Since we had already shown that dsRNA was within membrane-bound compartments, this left the question open as to whether the sites of viral RNA synthesis might be on the outside of these vesicles. DF1 cells were infected and treated with BrU and ActD as before, followed by IF labeling using either TX100 or digitonin permeabilization. As in previous experiments, the labeling pattern of newly synthesized viral RNA in TX100-permeabilized cells increased through the course of infection from smaller puncta to larger foci, mostly centered in perinuclear regions (Fig 3, top row). Through the course of infection, nsp12 signal was unaffected by digitonin permeabilization (Fig 3 and as before, in Fig 1). Strikingly however, much of the BrU signal at these timepoints was not detectable following digitonin permeabilization (Fig 3, bottom row). Although some newly synthesized viral RNA was found in the cytoplasm, this observation indicates that a large proportion of newly synthesized viral RNA is bound within a membrane. Overall viral RNA labeling followed a very similar staining pattern to dsRNA (as seen in Fig 1) and suggests that nascent viral RNA is located within the same membrane-bound compartments as dsRNA.

**Figure 3.**
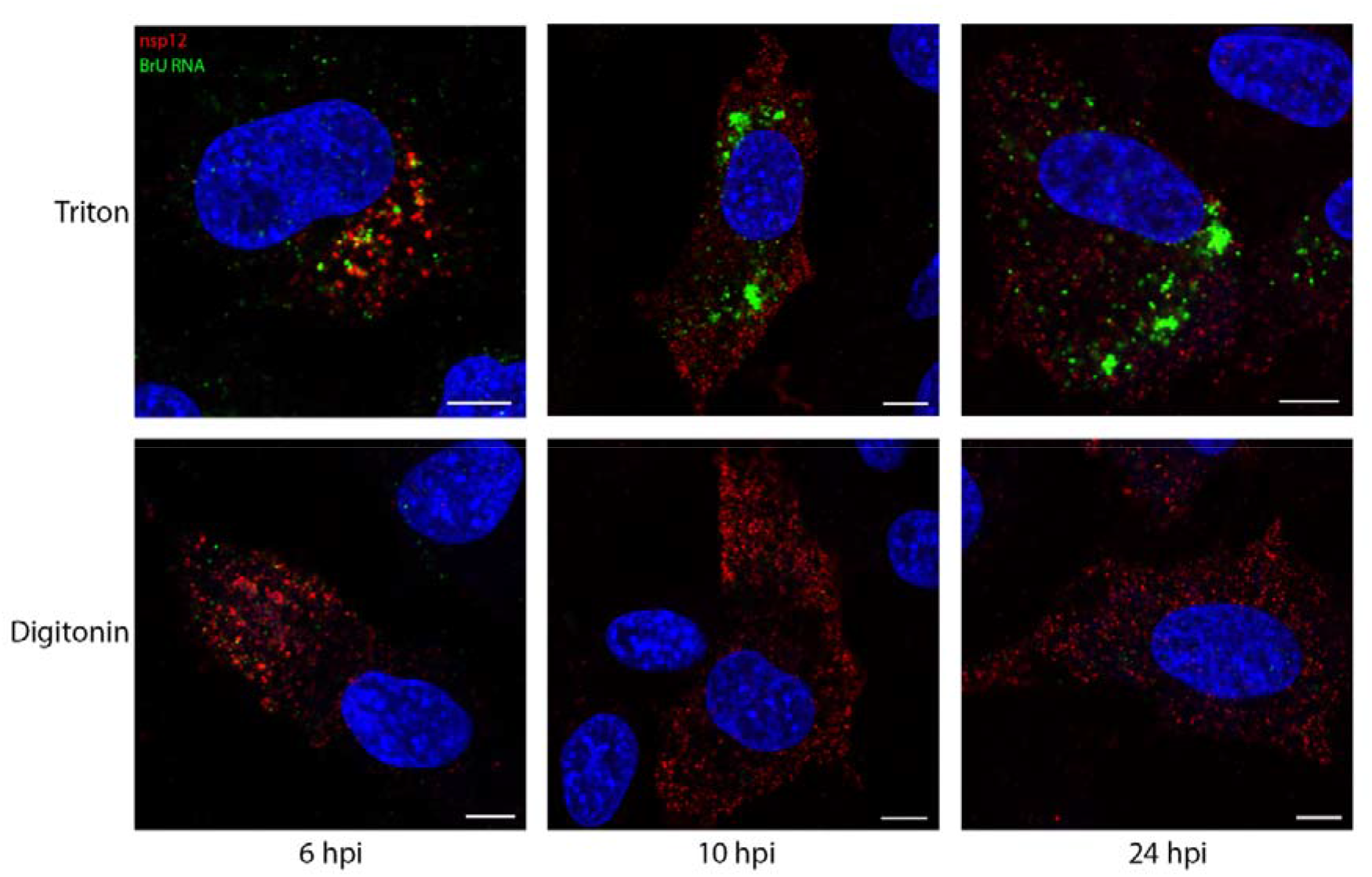
Viral RNA synthesis takes place in a membrane-bound compartment. DF1 cells were infected with IBV. 30 mins prior to fixation, cells were treated with BrU and ActD. Cells were fixed at the indicated times post infection. Cells were permeabilized with Triton X-100 (all membranes; top row) or digitonin (plasma membrane; bottom row) then labeled for nsp12 (red) and BrU (green), nuclei labeled with DAPI (blue). Scale bars represent 5 μm.

### Viral RNA is transported to the cytoplasm later in infection

To trace the fate of viral RNA synthesized early in infection and find out whether viral RNA produced within membrane-bound compartments is transported to the cytoplasm, cells were successively exposed to labeled uridine in the form of bromouridine (pulse) and then to unlabeled uridine (chase). In these pulse-chase experiments, viral RNA produced between 7-8 hpi was labeled with BrU as before, followed by a chase with excess unlabeled uridine until fixation at 24 hpi. The localization of viral RNA at 8 hpi was in cytoplasmic puncta, consistent with earlier observations (Fig 4a, left). When this viral RNA was chased to 24 hpi, it was found localized in large cytoplasmic puncta but also diffuse within the cytoplasm. (Fig 4a, right). To gain further information, cells were permeabilized with TX100 or digitonin as before. In control cells fixed at 8 hpi and permeabilized with TX100 or digitonin, labeling of the viral RNA was as before (Fig 4b, left). In the pulse-chased samples, the large foci of BrU-labeled RNA were no longer visible. However, the diffuse cytoplasmic signal was still detected (Fig 4b, right). Therefore, one pool of RNA was located within a membrane-bound compartment and a second was free in the cytoplasm. To investigate whether the cytoplasmic viral RNA might be in the process of being packaged, we looked to confirm that it colocalizes with viral structural proteins. Therefore, cells were labeled with an antibody specific for viral structural proteins (spike, membrane and nucleocapsid proteins). The colocalization between these markers (Fig 4c, top) indicates that the diffuse cytoplasmic staining pattern of BrU-labeled viral RNA is associated with structural proteins. In contrast, dsRNA was shown to colocalize with the large membrane-bound foci (Fig 4c, bottom).

**Figure 4.**
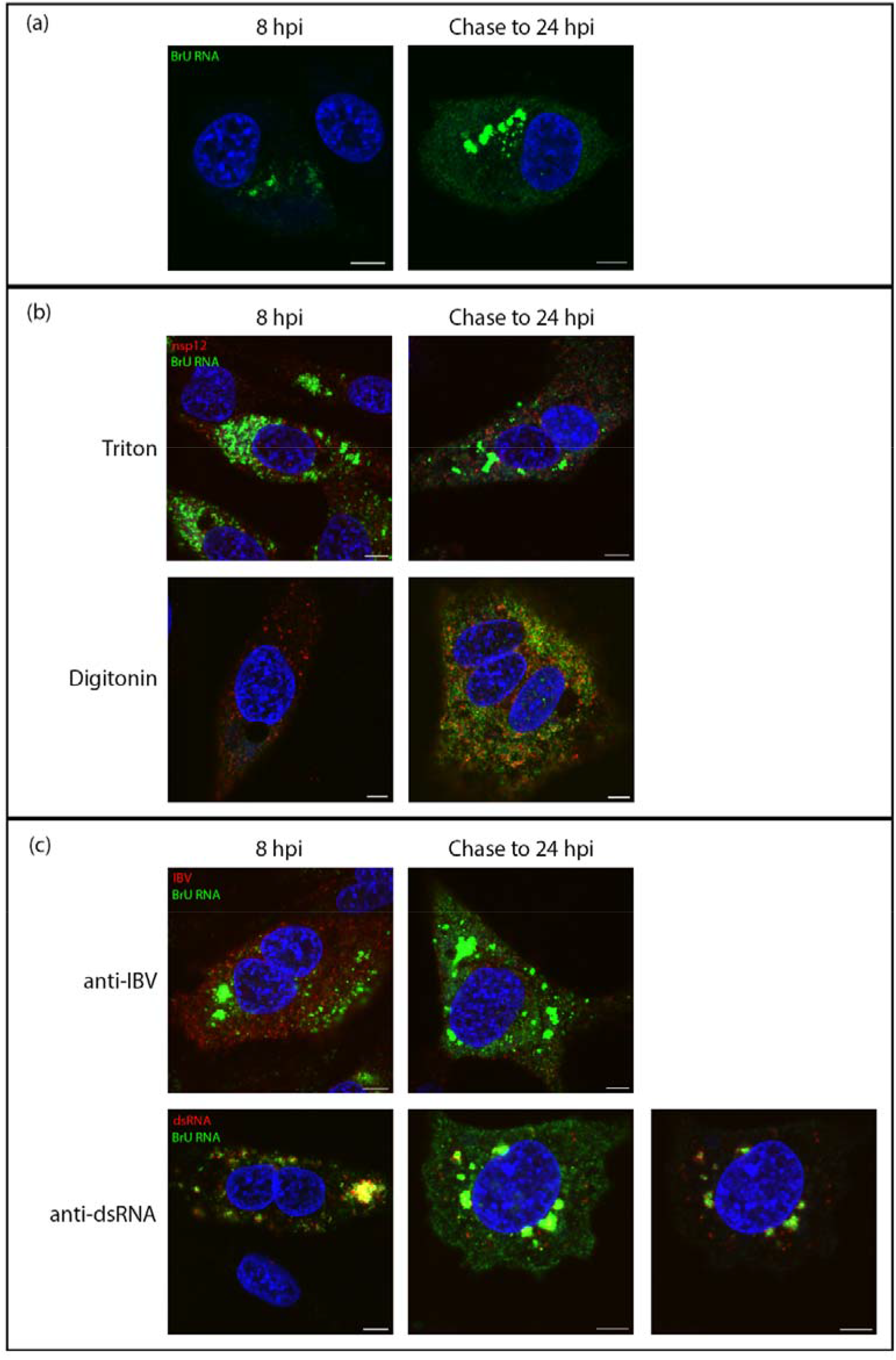
Viral RNA is transported to the cytoplasm later in infection. DF1 cells were infected with IBV. At 7 hpi cells were treated with BrU and ActD. At 8 hpi cells were either fixed (8 hpi) or chased with uridine until 24 hpi (chase to 24 hpi). Cells were labeled for BrU (green) and (b) nsp12, (c) IBV or (d) dsRNA (red), DAPI labeling nuclei (blue). Scale bars represent 5 μm.

### RNA synthesis by all genera of coronaviruses takes place in a membrane-bound compartment

While the structure of ROs has been shown to be conserved across all genera of CoVs, some morphological differences between the viruses do remain. Mainly, while convoluted membranes are found much more widely in alpha and beta CoV, they are found to a lesser extent in delta- and gamma-CoVs. In alpha- and beta-CoVs, the spherules were found associated with CM rather than zER and the majority of spherules were sealed compartments rather than remaining open to the cytosol as is the case in the majority of spherules in gamma-CoV infections [8]. We therefore sought to understand whether there might be any fundamental differences between the localization of sites of viral RNA synthesis in IBV compared with other CoV genera. Using one representative virus from each of the three other genera of CoVs (HCoV 229E (an alpha-CoV), SARS-CoV-2 (a beta-CoV) and PDCoV (a delta-CoV)), the location of nascent viral RNA labeled with BrU was assessed using TX100 and digitonin as before. Cells were labeled to detect BrU labeled nascent viral RNA or the viral nucleocapsid (N). While for each virus the N labeling (in red) was detected diffuse throughout the cytoplasm regardless of permeabilization method, BrU-labeled nascent viral RNA signal (in green) was contained within a membrane-bound compartment for HCoV 229E (Fig 5a), SARS-CoV-2 (Fig 5b) and PDCoV (Fig 5c). This is consistent with observations for IBV and demonstrates a conserved mechanism across the CoV family for viral RNA synthesis to be held within a membrane-bound compartment.

**Figure 5.**
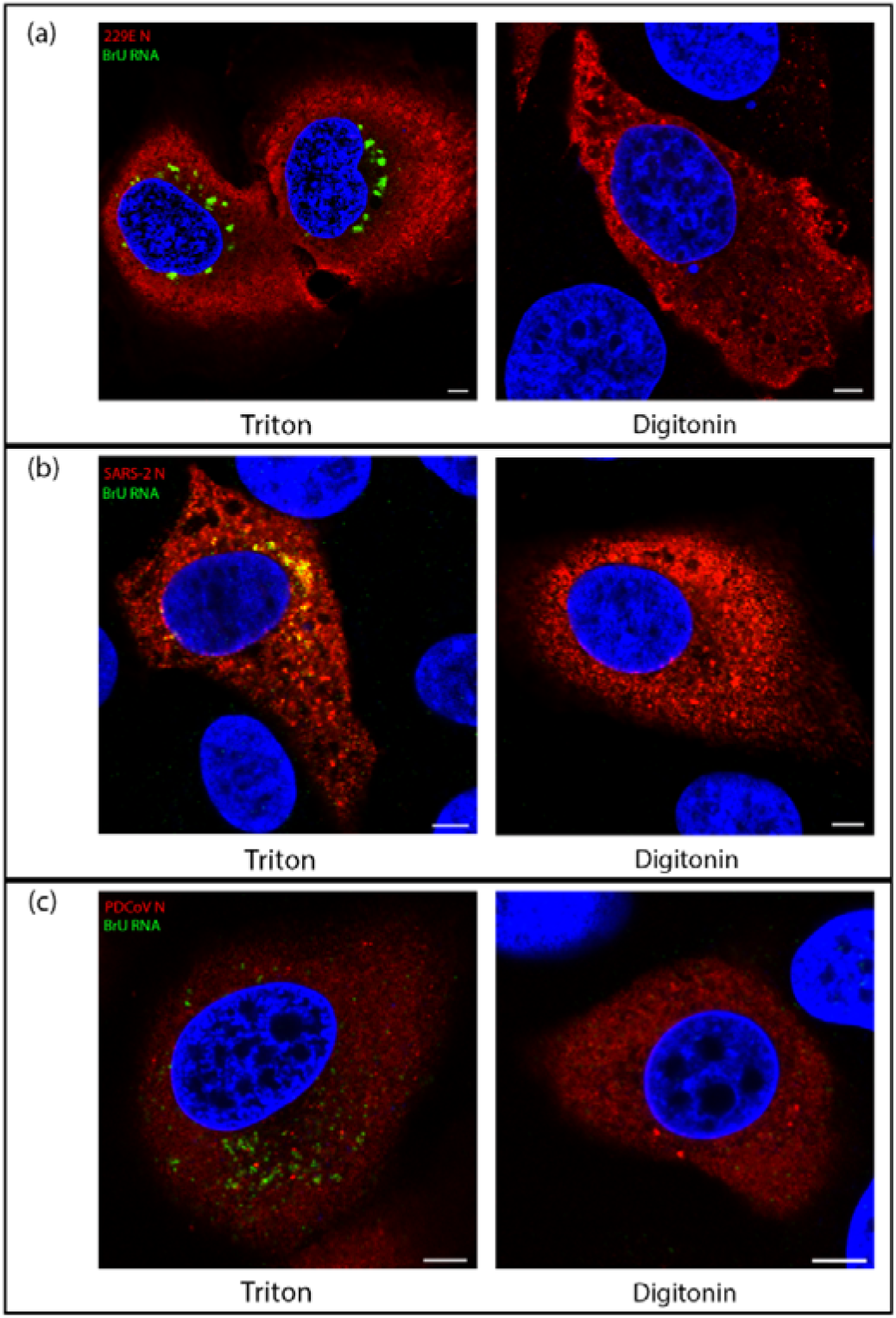
Viral RNA synthesis of diverse CoVs takes place within a membrane-bound compartment. (a) Huh7 cells were infected with HCoV 229E. At 7.5 hpi cells were treated with BrU and ActD for 30 min before fixation at 8 hpi. Cells were permeabilized with Triton X-100 (left) or digitonin (right) then labeled for BrU (green) and N (red), DAPI labeling the nuclei (blue). (b) VeroE6 cells were infected with SARS-CoV-2. At 5.5 hpi cells were treated with BrU and ActD for 30 min before fixation at 6 hpi. Cells were permeabilized with Triton X-100 (left) or digitonin (right) then labeled for BrU (green) and N (red), DAPI labeling the nuclei (blue). (c) LLC-PK1 cells were infected with PDCoV. At 5.5 hpi cells were treated with BrU and ActD for 30 min before fixation at 6 hpi. Cells were permeabilized with Triton X-100 (left) or digitonin (right) then labeled for BrU (green) and N (red), DAPI labeling the nuclei (blue). Scale bars represent 5 μm.

### Discussion

The formation of ROs is conserved across all +RNA viruses, an essential step in their life cycle. These ROs have long been thought to provide a site for viral RNA synthesis and in fact have been shown to do so for several viruses [8, 30, 31]. While these structures do vary between the different virus families, there are many similarities, the most basic of them being the energy required to rearrange cellular membranes to this extent. As obligate intracellular parasites, viruses are highly efficient and aim entirely to produce new generations of infectious particles as soon as possible. That these viruses induce these structures at all indicates that they are an important step in their life cycle [41, 43]. The gammacoronavirus, IBV induces the formation of DMVs, zER and spherules which pinch out from, but remain tethered to the zER. While IBV DMVs were recently identified as a site for viral RNA synthesis, no role has yet been found for spherules [8].

Previous studies have shown for other +RNA viruses such as SARS-CoV, FHV, and most recently in SARS-CoV-2 that dsRNA is found associated with DMVs [30, 31, 39, 40]. In the current study we have shown using different permeabilization methods, that dsRNA is found located within membrane-bound compartments. This is consistent with previous work [39] and suggests that dsRNA is also shielded within DMVs during IBV infection. Although a pore has recently been shown to be present in the DMV membrane [44], these are short lived/transient and have a diameter of 2-3nm at their narrowest point, which would not be large enough to allow entry of an antibody complex of ~30 nm. It is worth noting however, that our data here cannot exclude the possibility of dsRNA association with spherules. The IBV spherule neck measured 4-5 nm [37], which would also not be large enough to allow entry of an antibody complex. As dsRNA is a known target for intracellular pattern recognition receptors, it is likely that the virus aims to shield the dsRNA from detection within membrane-bound compartments such as DMVs. Indeed, activation of interferon (IFN) signaling is delayed following IBV infection and it was suggested that the IFN response then seen later in IBV infection could be due to dsRNA “leaking” from DMVs [51]. However, this now does not seem to be the case as data presented in the current work demonstrated that dsRNA was sealed within a membrane compartment at 24 hpi, a timepoint after which IFN signaling has been activated. Therefore, other mechanisms must exist to allow activation of IFN signaling later in IBV infection and these remain to be elucidated.

Finding sites of coronavirus RNA synthesis has been a topic of much research in recent years. Using biochemical methods *in vitro*, it has been shown that SARS-CoV RNA synthesis takes place inside membrane-bound compartments [38]. More recently, it was shown that DMVs are the site of viral RNA synthesis during MERS-CoV and IBV replication [8]. However, the methods used in that study could not definitively show whether RNA synthesis takes place on the interior or cytoplasmic face of DMV membranes. A subsequent study showed that DMVs contain pores within the membrane connecting the interior of the vesicle with the cytoplasm [44], providing a route by which RNA synthesized inside DMVs could exit for translation or packaging. Despite this, the location of viral RNA synthesis either inside or on the outside of DMVs remained to be confirmed. Significantly, here, by employing both BrU labeling of nascent RNA and different permeabilization methods, we have shown that sites of coronavirus RNA synthesis are membrane protected. Therefore, we have demonstrated conclusively that viral RNA synthesis in fact takes place *within* a membrane membrane-bound compartment. Moreover, we have confirmed that viral RNA synthesis by diverse CoVs from each of the four coronavirus genera, including recently identified SARS-CoV-2, takes place within a membrane-bound compartment. Although we cannot exclude that CoV RNA synthesis takes place associated with spherules, as MERS-CoV and IBV RNA synthesis has been shown to be associated with DMVs [8], we can infer here that RNA synthesis takes place *within* DMVs, and that this is conserved across the whole CoV family.

While it has previously been shown that newly synthesized IBV RNA location overlaps to some extent with the nsp14 [52], little else is known about the localization of viral proteins to sites of viral RNA synthesis. Several viral nsps are known to be involved in RO formation, including nsp3, nsp4 and nsp6 [39, 41, 43, 53–55] and nsps are known to localize to DMVs and other RO membranes [34, 39, 56]. Here, we have shown that viral RNA is associated with dsRNA but not with nsp12. While it was not surprising that as a replicative intermediate, dsRNA was found in close association with sites of viral RNA synthesis, the finding that nsp12 is not near these sites is somewhat surprising. However, this is likely to be because the antibody is unable to bind to nsp12 assembled within the replication/transcription complex (RTC) and actively involved in RNA synthesis. Nsp8 has been shown to interact with both the N- and C-terminal ends of nsp12 [57] which could very likely render the antigenic sites within nsp12 inaccessible to something as large as an antibody. Nsp12 also interacts directly with other viral proteins. MHV nsp12 interacts with nsp15 [58] and possibly with nsp5, nsp8 and nsp9 [59]. SARS-CoV nsp12 has been found in complex with nsp7, nsp8 and nsp14 [60]. Therefore, it is very likely that the nsp12 labeling detected here represents nsp12 that is not located within RTCs or is located within RTCs not actively involved in RNA synthesis. The amount of nsp12 signal detected that is presumably not located within active RTCs is perhaps rather surprising. However, several nsps from other CoVs are known to localize to both DMVs and other RO membranes [34, 39, 56]. The role of non-RTC associated nsp12 during virus replication remains to be determined.

Following characterization of the location of nascent RNA throughout infection, changes in the location of this RNA as infection progressed were studied. Using a pulse-chase approach, RNA synthesized between 7-8 hpi was visualized at 24 hpi. This demonstrated that by 24 hpi, viral RNA that was produced between 7-8 hpi showed two separate pools. The first remained in large, membrane-bound foci. These large accumulations of labeled viral RNA continued to associate with dsRNA within these compartments. The second pool of viral RNA produced between 7-8 hpi and tracked to 24 hpi had been exported into the cytoplasm. This pool of RNA is associated largely with structural proteins, presumably bound by N, or as it was being packaged into new virions and is consistent with the recently characterized pore in the DMV membrane [44]. These observations are consistent with a model whereby newly synthesized positive sense RNAs are exported out of the DMV to be translated or packaged, while positive and negative sense RNA templates remain within the DMVs for further rounds of RNA synthesis.

The CoV family contains many pathogens of animal and human interest. The RO of CoVs from all four genera have been confirmed previously to comprise DMVs and double membrane spherules. Here we investigated the site of viral RNA synthesis of one virus from each CoV genus. In all viruses we investigated (HCoV 229E, SARS-CoV-2, IBV and PDCoV) the site of viral RNA synthesis was bound within a membrane, consistent with being located on the interior of DMVs [8, 44]. The role of CoV induced convoluted membranes, zippered ER and double membrane spherules remains elusive. However, knowing that all CoVs synthesize viral RNA within a membrane-bound compartment is a significant step in understanding the replication of this important virus family.

## Supplementary Materials

The following are available online at www.mdpi.com/xxx/s1, Figure S1: Cellular transcription is inhibited with Actinomycin D treatment, Figure S2: Sites of viral RNA synthesis are associated with dsRNA but do not colocalize with nsp12., Video S3: Sites of viral RNA synthesis do not colocalize with nsp12; Video S4: Sites of viral RNA synthesis are associated with dsRNA.

## Author Contributions

Conceptualization, H.J.M. and P.C.H. methodology, N.D., J.S., P.C.H. and H.J.M.; validation, N.D.; formal analysis, N.D.; investigation, N.D.; resources, N.D., P.C.H. and H.J.M.; data curation, N.D.; writing—original draft preparation, N.D. and H.J.M.; writing—review and editing, N.D., J.S., P.C.H., and H.J.M.; visualization, N.D.; supervision, H.J.M.; project administration, H.J.M.; funding acquisition, H.J.M. All authors have read and agreed to the published version of the manuscript.

## Acknowledgments

The authors would like to thank Prof Linda Saif, Ohio State University, for kindly providing porcine deltacoronavirus for use in this work. We would also like to thank Prof Miles Carroll, UK Health Security Agency, for kindly sharing SARS-CoV-2, hCov-19/England/02/2020.

