## Supplementary information for "Coronavirus RNA synthesis takes place within membrane-bound sites"

### Materials & methods:

Supp 1: DF1 cells seeded onto glass coverslips were infected or mock-infected as in 2.1. Cells were then treated with 2 mM bromouridine (BrU; Sigma Aldrich) and 15 µM actinomycin D (ActD; Sigma Aldrich) or carrier control for 30 min. Cells were washed in PBS, fixed in RNase-free paraformaldehyde (pfm), then labeled. Cells were permeabilized in 0.1% Triton X-100 in PBS for 15 min, then incubated in blocking buffer (0.1% fish gelatin [Sigma Aldrich] in PBS) for 1 h. Primary antibodies specific for bromouridine (Table 1) were diluted in blocking buffer and incubated on cells for 1 h. After washing, Alexa Fluor secondary antibody (Invitrogen) in blocking buffer was incubated on cells for 1 h, followed by washing, labeling of nuclei using 4',6-diamidino-2-phenylindole (DAPI; Sigma Aldrich) and mounting onto glass slides with Vectashield (Vector Labs, Peterborough). For labeling of BrU samples, in order to prevent loss of the BrU signal, immunofluorescence (IF) labeling was promptly carried out in an RNase-free environment in the presence of RNasin at 0.133 U/ml (Promega, Southampton; [49]). Cells were visualized using a Leica CLSM SP5, and images assembled using Adobe Photoshop.

Supp 2, 3 & 4: DF1 cells seeded onto glass coverslips were infected, treated and labeled as in 2.1-2.3. Cells were visualized using a Leica CLSM SP8 microscope (Leica Microsystems, Milton Keynes, UK). For 3D reconstructions, z-stacks covering whole cells were collected and the data processed into models using Imaris software 9.2 (Bitplane, Oxford Instruments, UK). Images were assembled using Adobe Photoshop.

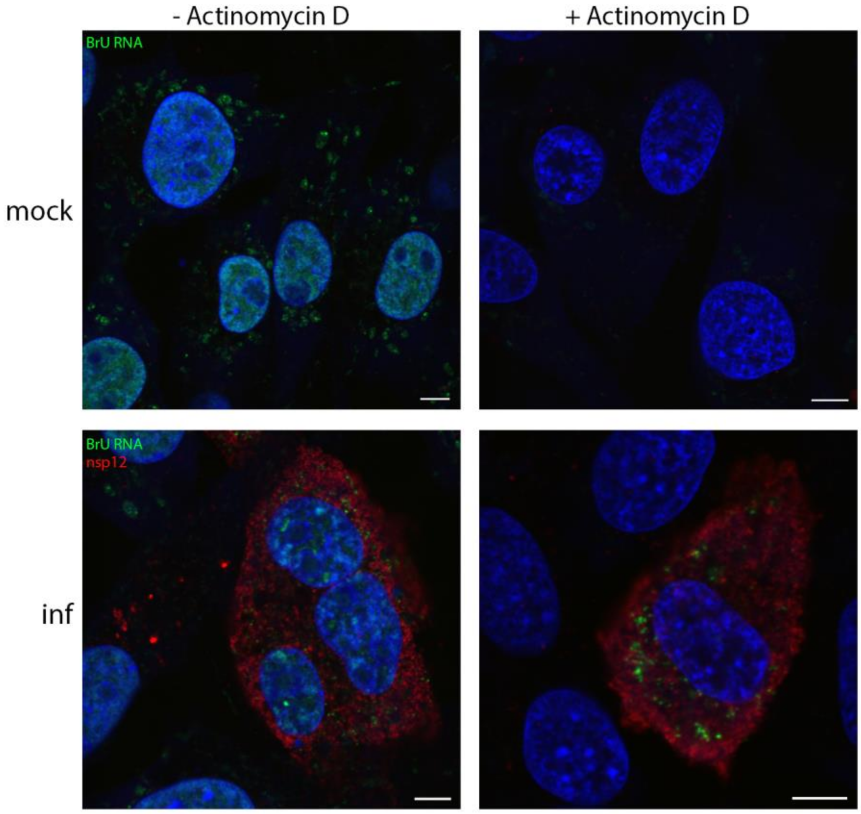


**Figure S1. Cellular transcription is inhibited with Actinomycin D treatment**
DF1 cells were infected or mock-infected, then treated with BrU and ActD (+ ActD) or carrier control (-ActD) for 30 min before fixation. They were then labeled for BrU (green), nsp12 (red) and DAPI (labeling the nuclei; blue). Scale bars represent 5 µm.

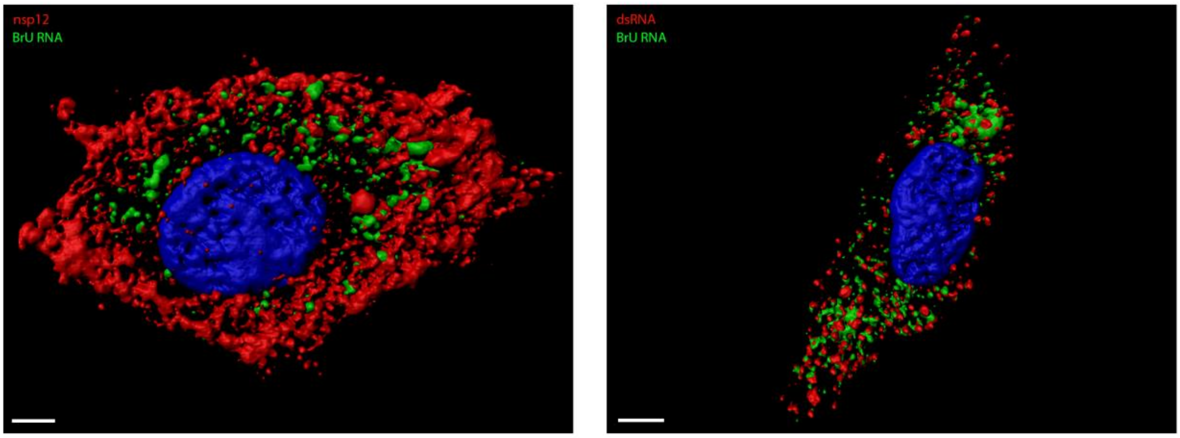


**Figure S2. Sites of viral RNA synthesis are associated with dsRNA but do not colocalize with nsp12.** DF1 cells were infected with IBV and 30 mins prior to fixation treated with BrU and ActD. Cells were fixed at 10 hpi and labeled with antibodies against BrU (green) and nsp12 (red; left) or dsRNA (red; right). Nuclei are labeled with DAPI (blue). Scale bars represent 5 μm. A z-stack was captured, then reconstructed using Imaris software 9.2.
